# Matched three-dimensional organoids and two-dimensional cell lines of melanoma brain metastases mirror response to targeted molecular therapy

**DOI:** 10.1101/2024.01.18.576318

**Authors:** William H. Hicks, Lauren C. Gattie, Jeffrey I Traylor, Diwakar Davar, Yana G. Najjar, Timothy E. Richardson, Samuel K. McBrayer, Kalil G. Abdullah

## Abstract

**Purpose:** Despite significant advances in the treatment paradigm for patients with metastatic melanoma, melanoma brain metastasis (MBM) continues to represent a significant treatment challenge. The study of MBM is limited, in part, by shortcomings in existing preclinical models. Surgically eXplanted Organoids (SXOs) are ex vivo, three-dimensional cultures prepared from primary tissue samples with minimal processing that recapitulate genotypic and phenotypic features of parent tumors and are grown without artificial extracellular scaffolding. We aimed to develop the first matched patient-derived SXO and cell line models of MBM to investigate responses to targeted therapy.

**Methods:** MBM SXOs were created by a novel protocol incorporating techniques for establishing glioma and cutaneous melanoma organoids. A BRAF^V600K^-mutant and BRAF-wildtype MBM sample were collected directly from the operating room for downstream experiments. Organoids were cultured in an optimized culture medium without an artificial extracellular scaffold. Concurrently, matched patient-derived cell lines were created. Drug screens were conducted to assess treatment response in SXOs and cell lines.

**Results:** Organoid growth was observed within 3-4 weeks, and MBM SXOs retained histological features of the parent tissue, including pleomorphic epithelioid cells with abundant cytoplasm, large nuclei, focal melanin accumulation, and strong SOX10 positivity. After sufficient growth, organoids could be manually parcellated to increase the number of replicates. Matched SXOs and cell lines demonstrated sensitivity to BRAF and MEK inhibitors.

**Conclusion:** Here, we describe the creation of a scaffold-free organoid model of MBM. Further study using SXOs may improve the translational relevance of preclinical studies and enable the study of the metastatic melanoma tumor microenvironment.

## Introduction

Melanoma brain metastasis (MBM) represents a significant treatment challenge [1]. While combinatorial systemic targeted therapy in some subtypes has shown reasonable control rates in extracranial tumor burden, this treatment regimen has produced a less durable response in progressive central nervous system (CNS) disease [2-4]. The recent CheckMate 204 trial demonstrated concordant intracranial and extracranial benefits with combined immune checkpoint inhibitors; however, further study is needed to assess optimal therapy regimens, particularly in symptomatic MBM [5, 6]. The decreased effectiveness of systemic targeted therapy to brain metastases has been thought to be related to poor blood-brain barrier penetration, the unique brain-tumor interface, and immune considerations of the brain-tumor immune microenvironment (TIME) [7-9]. While these interactions are critical to investigate, existing model systems have limited our ability to evaluate the response of MBM to immunotherapy in the laboratory.

Two-dimensional melanoma cell lines are essential for high-throughput, in vitro drug sensitivity studies; however, these systems do not reflect the complex cellular heterogeneity of human metastatic brain tumors [10]. Patient-derived xenograft models enable studies of drug responses in a relevant cellular milieu but are intrinsically low-throughput, carry long latency times, and the strains used for xenotransplantation limit evaluation of the TIME [11, 12]. Patient-derived three-dimensional organoids have emerged as an attractive system for modeling the heterogenous microenvironment of advanced cancers [13]. Traditional approaches to organoid modeling have used isogenic cell lines or enzymatically dissociated patient-derived tumor samples [14]. However, these techniques often simultaneously rely on the supplementation of exogenous growth factors and sample engrafting in an artificial extracellular matrix (ECM), such as Matrigel, Geltrex, or collagen gel scaffolds [13, 15]. Additionally, their clonal nature limits elements of the TIME that may drive treatment response or resistance.

Recently, our group demonstrated an optimized and efficient method to produce Surgically eXplanted Organoids (SXOs) of low-grade glioma from resected tumor samples that faithfully maintain the histology, genetics, and cellular composition of the parent tissue [16]. These SXOs can be established at an almost 90% success rate from both low- and high-grade gliomas and model the differential sensitivities to drugs that selectively target *IDH*-mutant gliomas [16, 17]. Notably, this approach to producing SXOs does not require an artificial ECM, thus providing a more faithful and physiologically relevant ex vivo model system. Recently, a minimally processed technique was used to generate patient-derived organoids from cutaneous melanoma samples [18, 19]. While this technique has clear advantages over protocols that dissociate tissues to single or near-single cell suspensions, it requires engraftment into an artificial ECM, is derived from a clonal progenitor cell, and exists in a system that lacks crucial elements of the TIME. To create an SXO model of MBM, we sought to adapt these techniques to incorporate elements of the glioma SXO model approach with variations in media concentrations to facilitate the growth of these metastatic lesions [16, 18, 19].

Herein, we detail the creation of three-dimensional, scaffold-free SXOs of MBM directly from primary patient tumor samples (**Fig. 1**). Additionally, we show that the MBM SXOs retain the histopathologic characteristics of the parent tissue and mirror predicted sensitivities to targeted therapeutics. To our knowledge, this is the first report of a scaffold-free organoid model of melanoma and is the first organoid model of MBM. These findings suggest that MBM SXOs may serve as a novel, physiologically relevant model system for therapeutic screening and translational studies of this disease.

**Fig. 1.**
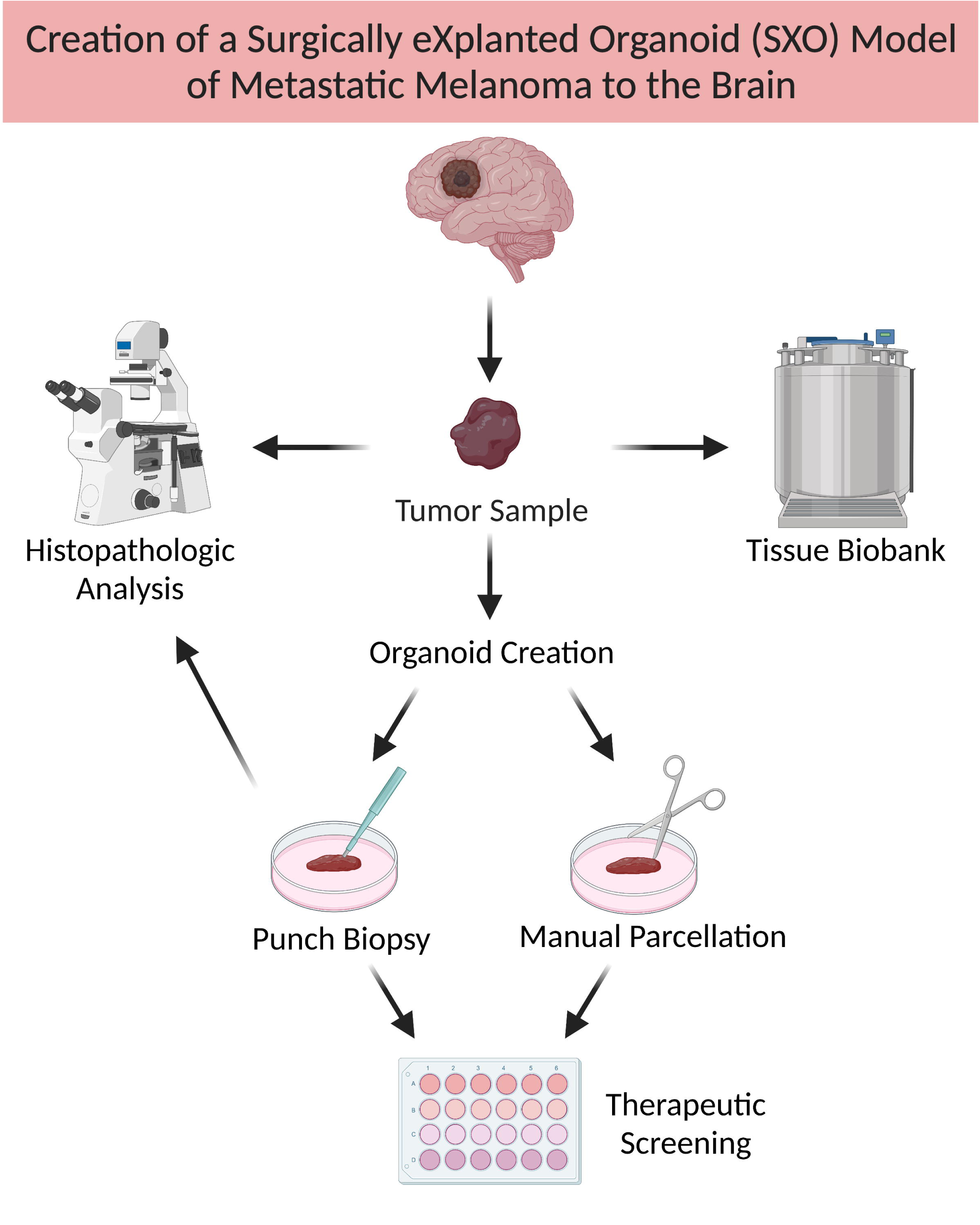
Overview of the creation of Surgically Explanted Organoids (SXOs) of Melanoma Brain Metastases (MBM). Tissue samples of MBM were collected from the operating room, manually parcellated, and cultured in optimized Melanoma Organoid Complete Media. SXOs reached maturity 2-4 weeks after explant and were cultured for at least four weeks before use in downstream experiments. Additionally, primary tissue and derivative organoids were cryopreserved for future analyses. Organoids were randomized to all treatment groups. Figure created with BioRender.

## Materials and Methods

### Human specimens

All patient tissue and blood samples were collected following ethical and technical guidelines on the use of human samples for biomedical research at the University of Pittsburgh and the University of Pittsburgh Medical Center after informed consent was obtained from patients under the Institutional Review Board Protocol (19080321). The study was conducted according to the principles of the Declaration of Helsinki.

### Organoid creation and culture in melanoma complete media

Tumor tissue was collected and processed as previously described [16]. Freshly resected tumor tissue was collected directly from the operating room, suspended in ice-cold Hibernate A (BrainBits HA), and brought to the lab on ice for processing within 30 minutes of resection. Tumor pieces were submerged in RBC Lysis Buffer (ThermoFisher 00433357) and incubated on a rocker at room temperature for 10 minutes. Then, the RBC Lysis Buffer was aspirated, and the tumor pieces were washed with Hibernate A solution containing Glutamax (2 mM, ThermoFisher 35050061), Penicillin/Streptomycin (100 U/mL and 100 μg/mL, respectively, ThermoFisher 15140122), and Amphotericin B (0.25 μg/mL, Gemini Bio-Products 400104). Tissues were transferred to a sterile culture dish, manually parcellated using a 1.0 mm biopsy punch (ThermoFisher 12460401), and suspended in 1 mL of Melanoma Organoid Complete Media (formula below) [18]. Each organoid in 1 mL of media was plated per well of a 24-well flat-bottom ultra-low adherence plate (Corning 3473). Plates were incubated in a humidified incubator at 37°C, 5% CO_2_, and 21% O_2_ under continuous orbital rotation at 120 rpm. SXO media was replaced every other day, with 75-90% of media being replaced. Melanoma Organoid Complete Media formula [18]: 250 mL DMEM (Sigma D6429), 125 mL Nutrient Mixture Ham’s F12 (Sigma N4888), 125 mL MCDB 105 Medium (Cell Applications 117-500), 50 mL fetal bovine serum (Gibco 26140079), 5 mL Glutamax (2 mM, ThermoFisher 35050061), 5 mL Penicillin/Streptomycin (100 U/mL and 100 μg/mL, respectively, ThermoFisher 15140122), 10 mL B-27 supplement (Gibco 17504044), 1 mL Normocin (Invivogen ant-nr-1). Melanoma Organoid Complete Media stocks were used up to 1 month after preparation.

### Human melanoma brain metastasis cell generation

The MBM739 cell line was generated from a human melanoma brain metastasis (UPMC739). The Human Tumor Dissociation Kit (Miltenyi Biotec 130095929) was used per the manufacturer’s protocol to generate a single-cell suspension from the tumor sample. After dissociation, cells were cultured in DMEM (Sigma D6546) supplemented with 10% fetal bovine serum, 5 mL Glutamax (2 mM, ThermoFisher 35050061), and 5 mL Penicillin/Streptomycin (100 U/mL and 100 μg/mL, respectively, ThermoFisher 15140122) at 37°C, 5% CO_2_, and 21% O_2_. Cells were cultured in adherent culture dishes and passaged 1-2 times per week with 0.25% trypsin.

### Histopathology and immunohistochemistry

SXOs and primary tissue samples were fixed in 10% formalin for 1 hour and then washed and resuspended in 70% ethanol. After the alcohol wash, SXOs were transferred to the cap of a 1.5 mL cryotube, resuspended in 0.5% agarose gel, and allowed to solidify overnight at 4°C. The following morning, the agarose-organoid molds were transferred to a cassette, dehydrated in 10% formalin, 70% ethanol three times for 30 minutes, 90% ethanol twice for 30 minutes, 100% ethanol three times for 30 minutes, xylene three times for 20 minutes, and embedded in paraffin. Paraffin blocks were sectioned at 4-um thickness. For hematoxylin and eosin (H&E) staining, slides were deparaffinized in xylene (9 min), 100% ethanol (3 min), 95% ethanol (1 min), and stained with hematoxylin (20 sec). Slides were rinsed in water, soaked in clarifier (40 sec), washed in water, and then bluing agent (20 sec). Slides were then rinsed, stained with eosin (20 sec), followed by serial incubation in 100% ethanol (3 min) and xylene (3 min), and then mounted for microscopic examination. Immunohistochemistry (IHC) staining was performed via the Cell Signaling Technology® citrate unmasking protocol. Antibodies used included an anti-human Sox10 rabbit antibody (1:100, Abcam ab227680) and an anti-human Gp100 mouse antibody (1:100, Abcam ab732, clone HMB45).

### Cell viability assay

Cell viability assays were performed by seeding 5,000 cells/well in a 96-well opaque walled plate. Replicates were then treated with the BRAF inhibitor, dabrafenib (Sellekchem S2807), MEK inhibitor, trametinib (Selleckchem S2673), or combined treatment. The CellTiter-Glo Luminescent Viability Assay (Promega G7571) was used to measure cell viability at 24, 48, and 72 hours following the manufacturer’s protocol. Luminescence was measured using the Tecan M1000 Pro Microplate Reader.

### Organoid viability assay

Organoids were treated with the respective drug diluted in Melanoma Complete Media. At the desired time point, SXO viability was assessed using the ReadyProbe Cell Viability Kit (Invitrogen R37610). Briefly, treated organoids were removed from the incubator, approximately 90% of the media was removed, and 500 μl fresh media was added. Then, 20 μL of the NucBlue/Hoechst and 20 μL of the propidium iodide dye were added. Organoids were then returned to the incubator under orbital rotation for 20 minutes. Fluorescent live cell imaging of the organoids was then captured on the Nikon CrestOptics X-Light V3 spinning disk confocal microscope at 40X magnification. An executable application was run within the Nikon NIS-Elements software to denoise and generate a three-dimensional maximal projection of the organoid. Organoid viability was quantified using the Nikon NIS-Elements software to generate the percent viability of live cells (Hoechst positive) relative to dead cells (Hoechst plus propidium iodide). Prior correlation of cellularity and viability between live cell imaging and traditional histopathology establishes the utility of this assay in assessing treatment response [20].

### Statistical analysis

SXOs were allocated to experiments randomly. All statistical tests were two-sided, where applicable. Student’s t-tests were used to assess the statistical significance of differences between groups. Statistical analyses were performed with GraphPad Prism (9.5.1.528, GraphPad Software, LLC) and included both descriptive statistics and tests of statistical significance. All data are plotted as mean ± standard deviation. For all tests, *p-*values less than 0.05 were considered statistically significant.

## Results

We created two unique models of MBM. Patient 1 (UPMC739) was a 54-year-old male with a 5-year history of BRAF^V600K^-mutant oligometastatic melanoma to the bowel and kidney who had undergone multiple extracranial resections as well as nivolumab and encorafenib/binimetinib treatments following multiple episodes of disease progression. He presented with new symptomatic multifocal brain metastases and underwent resection of a symptomatic left temporal lobe mass (**Fig. 2a**). Patient 2 (UPMC754) was a 62-year-old male with a 2-year history of BRAF-wildtype metastatic melanoma to the lungs who had undergone prior resection and treatment with nivolumab and duvelisib. He underwent resection of a symptomatic new large left cerebellar mass in the setting of multifocal MBM (**Fig. 2b**). Clinical pathologies were consistent with metastatic melanoma, denoted by epithelioid tumor cells with amphophilic cytoplasm, severe pleomorphism, prominent nucleoli, and variable (UPMC739) or absent (UPMC754) pigment staining. The tumor cells were positive for SOX10, gp100, and patchy MelanA by immunohistochemistry (IHC). From each tumor sample, we successfully generated SXOs and cultured them in Melanoma Organoid Complete Media [18] for four weeks before treatment studies. In parallel, we established a two-dimensional cell line, MBM739, from the primary tissue of UPMC739 using an available human tumor dissociation protocol. Cells were passaged weekly until capable of sufficient self-renewal for downstream assays.

**Fig. 2.**
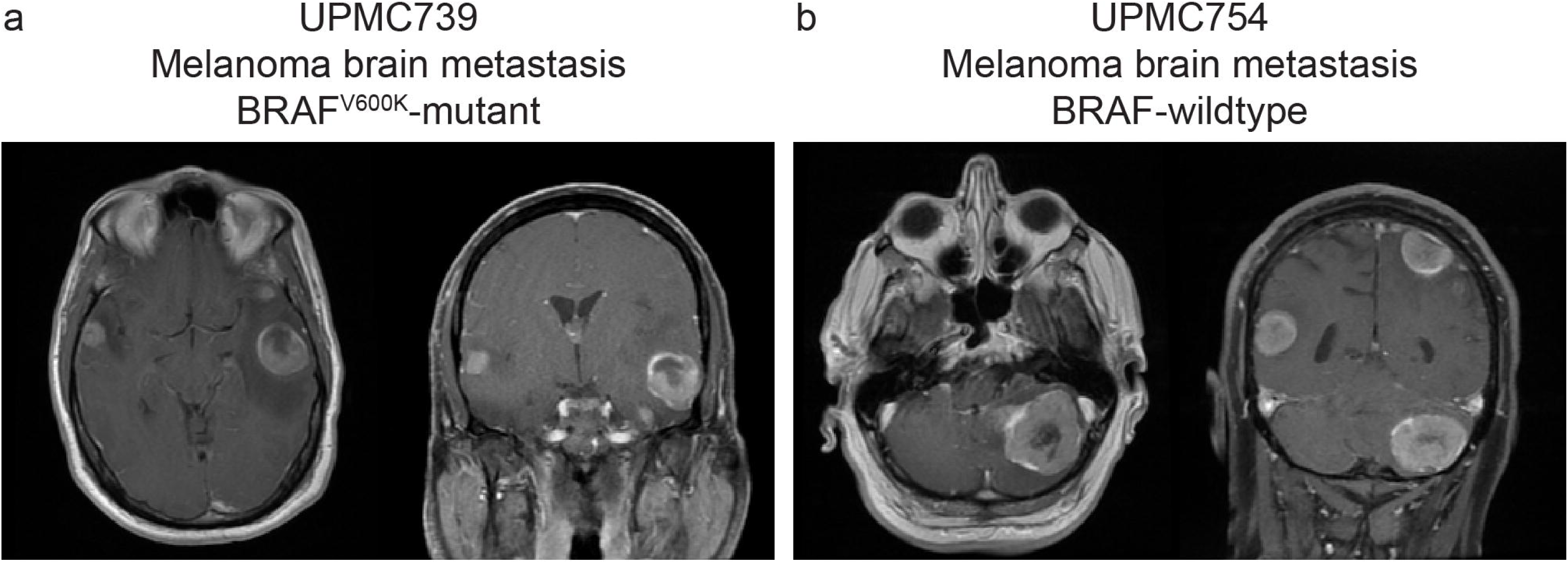
Preoperative axial and coronal T1 post-contrast magnetic resonance imaging (MRI) of (a) Patient 1 (UPMC739) revealing a large left temporal lobe melanoma metastasis, and (b) Patient 2 (UPMC754) revealing a large left cerebellar melanoma metastasis.

### MBM SXOs preserve histologic features of the parent tumor

We assessed SXO cytoarchitecture and viability with assistance from a board-certified neuropathologist (T.E.R.). We found that the SXO models recapitulated the histopathologic features of the parent tumor on hematoxylin and eosin (H&E) staining, with epithelioid cells and pleomorphic nuclei (**Fig. 3**). On visual inspection, the SXOs from UPMC739 were melanotic in appearance. In contrast, the SXOs from UPMC754 were tan and amelanotic, consistent with the degree of pigmentation present in the primary tumor histology. These pigmentation patterns were retained in culture, both grossly and on histopathology. Additionally, the IHC of both SXO models demonstrated diffuse positivity for SOX10, a reliable marker of multiple histologic subtypes of malignant melanoma [21], and UPMC739 was positive for gp100.

**Fig. 3.**
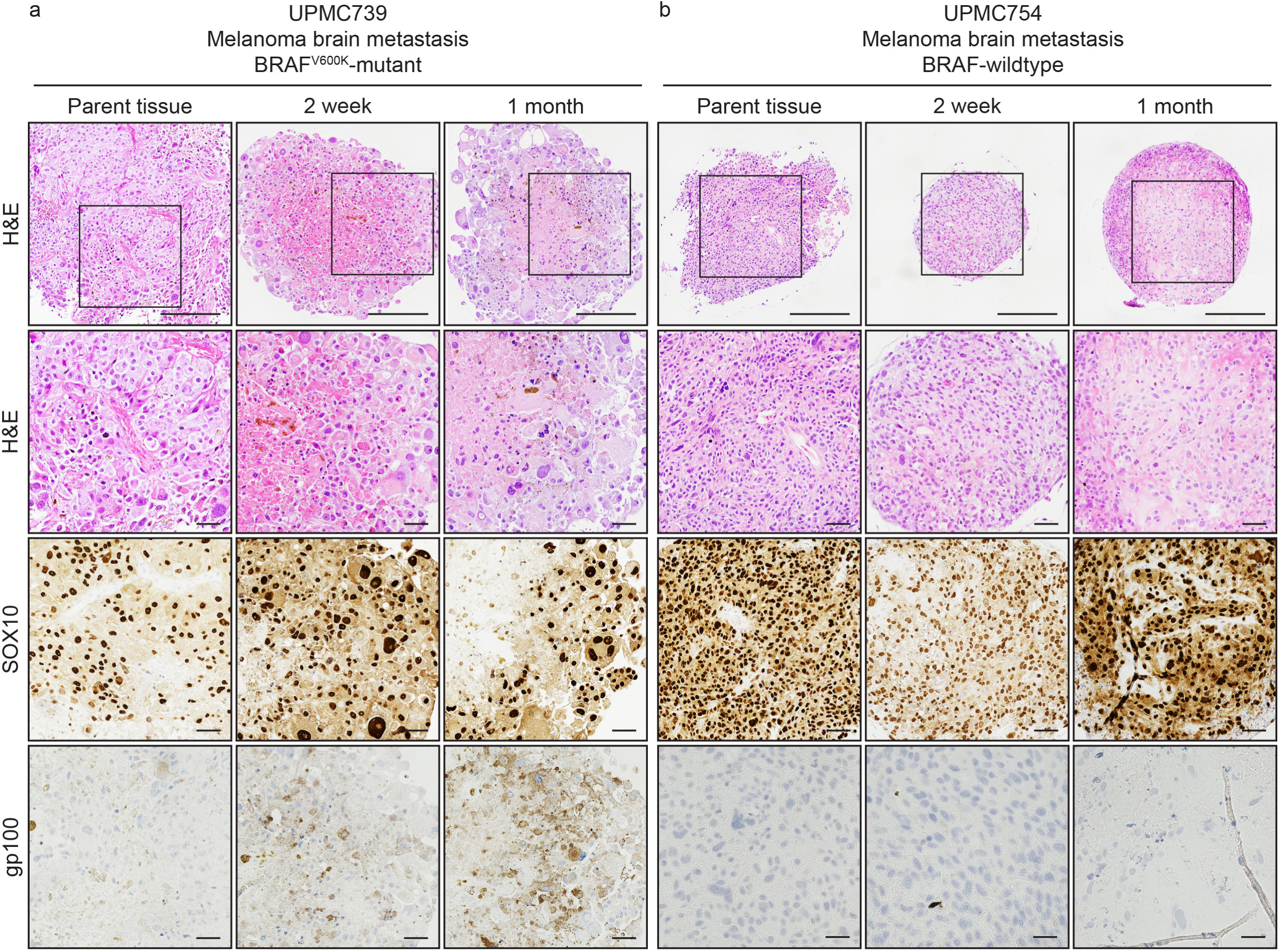
Surgically eXplanted Organoids (SXOs) of melanoma brain metastases retain the cellularity and histopathologic characteristics of the parent tissue. Explanted tissue and organoids were formalin-fixed and paraffin-embedded for histopathologic analysis at routine intervals, including at the time of surgery, two weeks post-resection, and one month post-resection. H&E slides demonstrate similar histological characteristics of SXOs to the parent tissue, including pleomorphic epithelioid cells with abundant cytoplasm, large nuclei, and focal (UPMC739) or absent (UPMC754) melanin accumulation. Immunohistochemistry shows that primary tissue and SXOs are diffusely SOX10 positive and gp100 positive (UPMC739). Scale bar = 125 μm and insets = 50 μm.

### Matched patient-derived cell line and surgically eXplanted organoids (SXOs) demonstrated sensitivity to dabrafenib and trametinib

To evaluate whether our patient-derived MBM models demonstrate predicted sensitivities to targeted molecular therapies, we treated the MBM739 cell line with dabrafenib (BRAF inhibitor), trametinib (MEK inhibitor), or combined BRAF-MEK inhibition with both agents (**Fig. 4a**). Using the doses determined from the cell line, we then administered the effective doses of each treatment to the MBM SXOs for 48 hours. We evaluated their response using an established method for live cell fluorescence imaging microscopy [20]. There was an appreciable increase in the propidium iodide (PI) signal with monotherapy treatment and a significant decrease in the organoid viability with dual-targeted molecular therapy (**Fig. 4b**). Further, Live-Dead analysis of the organoids treated with dabrafenib and trametinib alone showed a decrease in viability, however, combined BRAF-MEK inhibition resulted in a statistically significant decrease in MBM SXO viability (**Fig. 4c**)

**Fig. 4.**
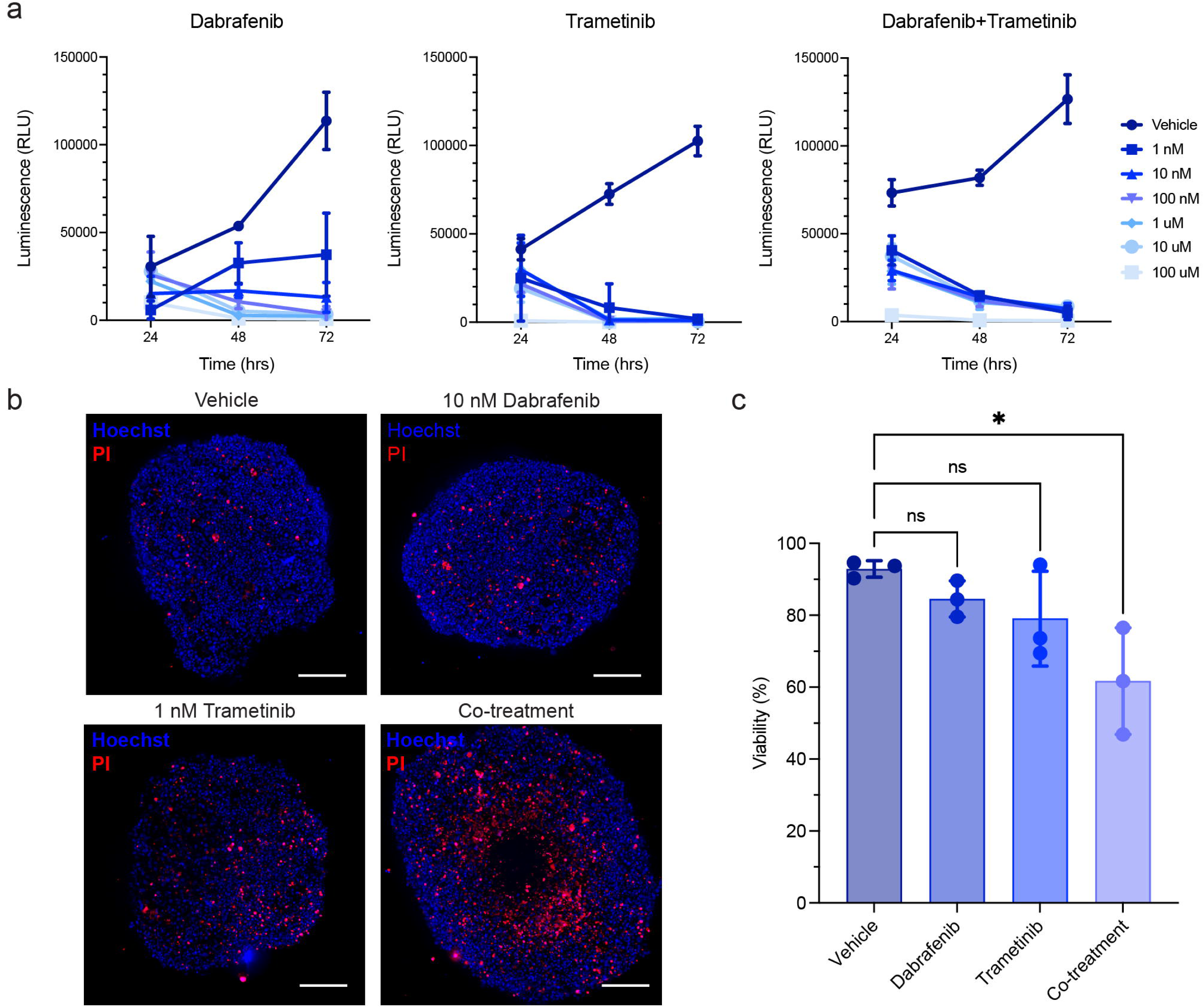
Matched patient-derived cell line and surgically eXplanted organoids (SXOs) of BRAF^V600K^-mutant metastatic melanoma to the brain demonstrate sensitivity to combined targeted therapy agents, dabrafenib and trametinib. (a) Decreased viability of the two-dimensional cell line MBM739, derived from UPMC739, in response to increasing dabrafenib, trametinib, or combined agents as assessed by the Cell-Titer Glo assay. (b) Fluorescent-based viability assay of MBM SXOs revealed maintenance of organoid structural integrity with increased cell death after 48 hours of treatment. (c) Quantification of SXO viability demonstrates the mild effect of monotherapy and a synergistic effect with dual-targeted therapy. Data are presented as means ± standard deviation; ns = not significant; * p<0.05. n = 3 for all treatment groups. Scale bar = 250 μm.

## Discussion

Here, we detail a protocol for a novel preclinical model of MBM and the first known scaffold-free organoid model of melanoma. Further, we have shown the ability to screen the sensitivity of MBM SXOs to targeted therapeutic agents. The generation of this model answers a prior unmet need for a physiologically relevant preclinical system to guide translational studies of MBM.

Melanoma is highly aggressive and carries a significant mortality rate due to its rapid proliferation, early and recurrent metastatic disease propensity, particularly to the CNS, and intratumor heterogeneity [22, 23]. While combination targeted molecular therapy and immunotherapy have shown success in treating extracranial disease burden, the degree of disease response for MBM has been more limited. While a recent study demonstrated that treatment of asymptomatic MBM with nivolumab plus ipilimumab (Checkmate 204, open-label phase 2 results) was feasible, there remain critical subsets of patients who either develop symptomatic MBM or who progress despite combinatorial therapy [6].

Alternatively, the outcomes in MBM may be attributable to the paucity of relevant model systems that faithfully and reliably model human metastatic melanoma to the brain [24]. Two-dimensional cell lines frequently lack intratumoral heterogeneity, traditional xenograft models limit the study of the TIME, and genetically engineered mouse models often produce oligoclonal tumors compared to the polyclonal heterogeneity seen in human melanoma, particularly in the case of metastatic disease [24]. Thus, the protocol outlined here may bridge a translational gap by providing a preclinical MBM model that is temporally related to the primary human tumor, has minimal ex vivo processing, retains the key histopathologic features of the primary tumor, and mirrors predicted drug sensitivities of the primary tumor based on known molecular profiles.

A downstream application of this model system is in performing preclinical therapeutic screening. In addition to being useful in testing sensitivity to targeted molecular therapies, as seen in the current study and our prior studies targeting *IDH*-mutant glioma [16, 17], we have shown that SXOs provide reliable treatment responses in multi-modality evaluations of targeted therapeutics [25, 26]. Our study demonstrated the utility of this model system in assessing treatment response to targeted agents against upregulated cell signaling pathways in BRAF-mutant MBM [27-29]. Patient-derived organoids have been used to perform comprehensive drug screens in extracranial platforms for breast, liver, and colorectal cancers, among others [13, 30, 31]. Notably, the minimally processed SXO model can be produced in a clinically relevant timeline [32]. As with the results from our prior studies, the cultured MBM SXOs reach maturity within 2-4 weeks of creation, in line with the timing for postoperative recovery, histopathologic analysis of biopsied or resected tissue, and genomic sequencing profiles to result. With clinical practice for most hematologic and solid malignancies centered around individual cancer’s genetic profile, a comprehensive preclinical drug screen of the patient’s tumor would be an additional tool to guide personalized oncology [33].

Beyond their initial applications in translational therapeutic screenings, SXOs offer a novel avenue to understand melanoma biology. Glioma SXOs have demonstrated preservation of the tumor, non-immune, and infiltrating immune cells, enabling the study of the tumor microenvironment [16, 34]. Particularly when studying melanoma, where systemic immunotherapy has shown significant efficacy, a model system to study these interactions ex vivo is critical. A recent study by Ou et al. created a scaffolded organoid model of extracranial malignant melanoma. They found they could successfully model the immunosuppressive microenvironment and sensitivity to immune checkpoint blockade in vivo [35]. Additionally, we have recently shown the ability to perform stable isotope tracing of glioma SXOs, providing another unique avenue to study tumor metabolism in metastatic cancer of the brain [36].

## Limitations

We acknowledge that our study is not without limitations. All replicates were derived from two patients and may not fully represent the generalizability of this protocol. Further, the sensitivity of the BRAF^V600K^-mutant MBM SXO to dabrafenib and trametinib may have been confounded by prior exposure to encorafenib and binimetinib, an alternative BRAF/MEK inhibitor combination. The number of SXO replicates is limited (n = 3 for histology and n = 3 for drug treatments). It should be noted that the generation and maintenance of SXOs is contingent on precise attention to culture media, and our study marries the culture needs of rapidly-derived tissue explants and melanoma-specific media, which was critical in the generation of these models. This is further supported by prior organoid studies in which different culture media conditions with the appropriate growth factors are required to create and propagate breast, lung, and brain cancer organoids [13, 16, 34]. Additionally, an expeditious relay between the operating room and the laboratory is essential, as these explants have shown high sensitivity to delays in culture conditions.

## Acknowledgments

Not applicable.

## Author Contributions

Conceptualization: K.G.A., S.K.M., W.H.H.; Methodology: W.H.H. and K.G.A.; Investigation: W.H.H., L.C.G., T.E.R.; Formal Analysis: W.H.H., L.C.G., T.E.R.; Writing – Original Draft: W.H.H., K.G.A.; Writing – Review & Editing: W.H.H., J.I.T., D.D., Y.G.N., L.C.G., S.K.M., and K.G.A.; Visualization: W.H.H., L.C.G.; Supervision: K.G.A. and S.K.M.; Funding Acquisition: K.G.A.

## Data Availability Statement

Requests for information, resources, and reagents should be directed to and will be fulfilled by the Lead Contact, William H. Hicks. Additional histopathology images will be shared by the lead contact upon request. Any additional information required to reanalyze the data reported in this paper is available from the lead contact upon request.

## Ethical Statement

All patient tissue and blood samples were collected following ethical and technical guidelines on the use of human samples for biomedical research at the University of Pittsburgh after informed consent was obtained from patients under the Institutional Review Board Protocol (19080321). The study was conducted according to the principles of the Declaration of Helsinki.

## References

1. Gutzmer R, Vordermark D, Hassel JC, Krex D, Wendl C, Schadendorf D, Sickmann T, Rieken S, Pukrop T, Höller C, Eigentler TK, Meier F (2020) Melanoma brain metastases – Interdisciplinary management recommendations 2020. Cancer Treat Rev 89: 102083 doi:10.1016/j.ctrv.2020.102083

2. Rishi A, Yu H-HM (2020) Current Treatment of Melanoma Brain Metastasis. Current Treatment Options in Oncology 21: 45 doi:10.1007/s11864-020-00733-z

3. Bander ED, Yuan M, Carnevale JA, Reiner AS, Panageas KS, Postow MA, Tabar V, Moss NS (2021) Melanoma brain metastasis presentation, treatment, and outcomes in the age of targeted and immunotherapies. Cancer 127: 2062–2073 doi:10.1002/cncr.33459

4. Davies MA, Saiag P, Robert C, Grob JJ, Flaherty KT, Arance A, Chiarion-Sileni V, Thomas L, Lesimple T, Mortier L, Moschos SJ, Hogg D, Márquez-Rodas I, Del Vecchio M, Lebbé C, Meyer N, Zhang Y, Huang Y, Mookerjee B, Long GV (2017) Dabrafenib plus trametinib in patients with BRAF(V600)-mutant melanoma brain metastases (COMBI-MB): a multicentre, multicohort, open-label, phase 2 trial. Lancet Oncol 18: 863–873 doi:10.1016/s1470-2045(17)30429-1

5. Tawbi HA, Forsyth PA, Algazi A, Hamid O, Hodi FS, Moschos SJ, Khushalani NI, Lewis K, Lao CD, Postow MA, Atkins MB, Ernstoff MS, Reardon DA, Puzanov I, Kudchadkar RR, Thomas RP, Tarhini A, Pavlick AC, Jiang J, Avila A, Demelo S, Margolin K (2018) Combined Nivolumab and Ipilimumab in Melanoma Metastatic to the Brain. N Engl J Med 379: 722–730 doi:10.1056/NEJMoa1805453

6. Tawbi HA, Forsyth PA, Hodi FS, Algazi AP, Hamid O, Lao CD, Moschos SJ, Atkins MB, Lewis K, Postow MA, Thomas RP, Glaspy J, Jang S, Khushalani NI, Pavlick AC, Ernstoff MS, Reardon DA, Kudchadkar R, Tarhini A, Chung C, Ritchings C, Durani P, Askelson M, Puzanov I, Margolin KA (2021) Long-term outcomes of patients with active melanoma brain metastases treated with combination nivolumab plus ipilimumab (CheckMate 204): final results of an open-label, multicentre, phase 2 study. The Lancet Oncology 22: 1692–1704 doi:10.1016/S1470-2045(21)00545-3

7. Murrell J, Board R (2013) The use of systemic therapies for the treatment of brain metastases in metastatic melanoma: opportunities and unanswered questions. Cancer Treat Rev 39: 833–838 doi:10.1016/j.ctrv.2013.06.004

8. Makawita S, Tawbi HA (2021) Nonsurgical Management of Melanoma Brain Metastasis: Current Therapeutics, Challenges, and Strategies for Progress. Am Soc Clin Oncol Educ Book 41: 79–90 doi:10.1200/edbk_321137

9. Rulli E, Legramandi L, Salvati L, Mandala M (2019) The impact of targeted therapies and immunotherapy in melanoma brain metastases: A systematic review and meta-analysis. Cancer 125: 3776–3789 doi:10.1002/cncr.32375

10. Zhou S, Lu J, Liu S, Shao J, Liu Z, Li J, Xiao Wa (2023) Role of the tumor microenvironment in malignant melanoma organoids during the development and metastasis of tumors. Frontiers in Cell and Developmental Biology 11 doi:10.3389/fcell.2023.1166916

11. Kircher DA, Silvis MR, Cho JH, Holmen SL (2016) Melanoma Brain Metastasis: Mechanisms, Models, and Medicine. International journal of molecular sciences 17 doi:10.3390/ijms17091468

12. Hicks WH, Bird CE, Traylor JI, Shi DD, El Ahmadieh TY, Richardson TE, McBrayer SK, Abdullah KG (2021) Contemporary Mouse Models in Glioma Research. Cells 10 doi:10.3390/cells10030712

13. Sachs N, de Ligt J, Kopper O, Gogola E, Bounova G, Weeber F, Balgobind AV, Wind K, Gracanin A, Begthel H, Korving J, van Boxtel R, Duarte AA, Lelieveld D, van Hoeck A, Ernst RF, Blokzijl F, Nijman IJ, Hoogstraat M, van de Ven M, Egan DA, Zinzalla V, Moll J, Boj SF, Voest EE, Wessels L, van Diest PJ, Rottenberg S, Vries RGJ, Cuppen E, Clevers H (2018) A Living Biobank of Breast Cancer Organoids Captures Disease Heterogeneity. Cell 172: 373–386.e310 10.1016/j.cell.2017.11.010

14. Pernik MN, Bird CE, Traylor JI, Shi DD, Richardson TE, McBrayer SK, Abdullah KG (2021) Patient-Derived Cancer Organoids for Precision Oncology Treatment. J Pers Med 11 doi:10.3390/jpm11050423

15. Kim M, Mun H, Sung CO, Cho EJ, Jeon HJ, Chun SM, Jung DJ, Shin TH, Jeong GS, Kim DK, Choi EK, Jeong SY, Taylor AM, Jain S, Meyerson M, Jang SJ (2019) Patient-derived lung cancer organoids as in vitro cancer models for therapeutic screening. Nat Commun 10: 3991 doi:10.1038/s41467-019-11867-6

16. Abdullah KG, Bird CE, Buehler JD, Gattie LC, Savani MR, Sternisha AC, Xiao Y, Levitt MM, Hicks WH, Li W, Ramirez DMO, Patel T, Garzon-Muvdi T, Barnett S, Zhang G, Ashley DM, Hatanpaa KJ, Richardson TE, McBrayer SK (2022) Establishment of patient-derived organoid models of lower-grade glioma. Neuro-oncology 24: 612–623 doi:10.1093/neuonc/noab273

17. Shi DD, Savani MR, Levitt MM, Wang AC, Endress JE, Bird CE, Buehler J, Stopka SA, Regan MS, Lin YF, Puliyappadamba VT, Gao W, Khanal J, Evans L, Lee JH, Guo L, Xiao Y, Xu M, Huang B, Jennings RB, Bonal DM, Martin-Sandoval MS, Dang T, Gattie LC, Cameron AB, Lee S, Asara JM, Kornblum HI, Mak TW, Looper RE, Nguyen QD, Signoretti S, Gradl S, Sutter A, Jeffers M, Janzer A, Lehrman MA, Zacharias LG, Mathews TP, Losman JA, Richardson TE, Cahill DP, DeBerardinis RJ, Ligon KL, Xu L, Ly P, Agar NYR, Abdullah KG, Harris IS, Kaelin WG, Jr., McBrayer SK (2022) De novo pyrimidine synthesis is a targetable vulnerability in IDH mutant glioma. Cancer cell 40: 939–956.e916 doi:10.1016/j.ccell.2022.07.011

18. Vilgelm AE, Bergdorf K, Wolf M, Bharti V, Shattuck-Brandt R, Blevins A, Jones C, Phifer C, Lee M, Lowe C, Hongo R, Boyd K, Netterville J, Rohde S, Idrees K, Bauer JA, Westover D, Reinfeld B, Baregamian N, Richmond A, Rathmell WK, Lee E, McDonald OG, Weiss VL (2020) Fine-Needle Aspiration-Based Patient-Derived Cancer Organoids. iScience 23: 101408 10.1016/j.isci.2020.101408

19. Phifer CJ, Bergdorf KN, Bechard ME, Vilgelm A, Baregamian N, McDonald OG, Lee E, Weiss VL (2021) Obtaining patient-derived cancer organoid cultures via fine-needle aspiration. STAR Protocols 2: 100220 10.1016/j.xpro.2020.100220

20. Buehler JD, Bird CE, Savani MR, Gattie LC, Hicks WH, Levitt MM, El Shami M, Hatanpaa KJ, Richardson TE, McBrayer SK, Abdullah KG (2022) Semi-Automated Computational Assessment of Cancer Organoid Viability Using Rapid Live-Cell Microscopy. Cancer Inform 21: 11769351221100754 doi:10.1177/11769351221100754

21. Willis BC, Johnson G, Wang J, Cohen C (2015) SOX10: a useful marker for identifying metastatic melanoma in sentinel lymph nodes. Appl Immunohistochem Mol Morphol 23: 109–112 doi:10.1097/pai.0000000000000097

22. Keung EZ, Gershenwald JE (2018) The eighth edition American Joint Committee on Cancer (AJCC) melanoma staging system: implications for melanoma treatment and care. Expert Rev Anticancer Ther 18: 775–784 doi:10.1080/14737140.2018.1489246

23. Shannan B, Perego M, Somasundaram R, Herlyn M (2016) Heterogeneity in Melanoma. Cancer Treat Res 167: 1–15 doi:10.1007/978-3-319-22539-5_1

24. Gregg RK (2021) Model Systems for the Study of Malignant Melanoma. In: Hargadon KM (ed) Melanoma: Methods and Protocols. Springer US, New York, NY, pp 1-21

25. Yu S, Wei S, Savani M, Lin X, Du K, Mender I, Siteni S, Vasilopoulos T, Reitman ZJ, Ku Y, Wu D, Liu H, Tian M, Chen Y, Labrie M, Charbonneau CM, Sugarman E, Bowie M, Hariharan S, Waitkus M, Jiang W, McLendon RE, Pan E, Khasraw M, Walsh KM, Lu Y, Herlyn M, Mills G, Herbig U, Wei Z, Keir ST, Flaherty K, Liu L, Wu K, Shay JW, Abdullah K, Zhang G, Ashley DM (2021) A Modified Nucleoside 6-Thio-2’-Deoxyguanosine Exhibits Antitumor Activity in Gliomas. Clinical cancer research: an official journal of the American Association for Cancer Research 27: 6800–6814 doi:10.1158/1078-0432.Ccr-21-0374

26. Wei S, Yin D, Yu S, Lin X, Savani MR, Du K, Ku Y, Wu D, Li S, Liu H, Tian M, Chen Y, Bowie M, Hariharan S, Waitkus M, Keir ST, Sugarman ET, Deek RA, Labrie M, Khasraw M, Lu Y, Mills GB, Herlyn M, Wu K, Liu L, Wei Z, Flaherty KT, Abdullah K, Zhang G, Ashley DM (2022) Antitumor Activity of a Mitochondrial-Targeted HSP90 Inhibitor in Gliomas. Clinical cancer research: an official journal of the American Association for Cancer Research 28: 2180–2195 doi:10.1158/1078-0432.Ccr-21-0833

27. Zhuang L, Lee CS, Scolyer RA, McCarthy SW, Palmer AA, Zhang XD, Thompson JF, Bron LP, Hersey P (2005) Activation of the extracellular signal regulated kinase (ERK) pathway in human melanoma. J Clin Pathol 58: 1163–1169 doi:10.1136/jcp.2005.025957

28. Haass NK, Sproesser K, Nguyen TK, Contractor R, Medina CA, Nathanson KL, Herlyn M, Smalley KS (2008) The mitogen-activated protein/extracellular signal-regulated kinase kinase inhibitor AZD6244 (ARRY-142886) induces growth arrest in melanoma cells and tumor regression when combined with docetaxel. Clinical cancer research: an official journal of the American Association for Cancer Research 14: 230–239 doi:10.1158/1078-0432.Ccr-07-1440

29. Bollag G, Hirth P, Tsai J, Zhang J, Ibrahim PN, Cho H, Spevak W, Zhang C, Zhang Y, Habets G, Burton EA, Wong B, Tsang G, West BL, Powell B, Shellooe R, Marimuthu A, Nguyen H, Zhang KYJ, Artis DR, Schlessinger J, Su F, Higgins B, Iyer R, D’Andrea K, Koehler A, Stumm M, Lin PS, Lee RJ, Grippo J, Puzanov I, Kim KB, Ribas A, McArthur GA, Sosman JA, Chapman PB, Flaherty KT, Xu X, Nathanson KL, Nolop K (2010) Clinical efficacy of a RAF inhibitor needs broad target blockade in BRAF-mutant melanoma. Nature 467: 596–599 doi:10.1038/nature09454

30. Ji S, Feng L, Fu Z, Wu G, Wu Y, Lin Y, Lu D, Song Y, Cui P, Yang Z, Sang C, Song G, Cai S, Li Y, Lin H, Zhang S, Wang X, Qiu S, Zhang X, Hua G, Li J, Zhou J, Dai Z, Wang X, Ding L, Wang P, Gao D, Zhang B, Rodriguez H, Fan J, Clevers H, Zhou H, Sun Y, Gao Q (2023) Pharmaco-proteogenomic characterization of liver cancer organoids for precision oncology. Science Translational Medicine 15: eadg3358 doi:10.1126/scitranslmed.adg3358

31. van de Wetering M, Francies Hayley E, Francis Joshua M, Bounova G, Iorio F, Pronk A, van Houdt W, van Gorp J, Taylor-Weiner A, Kester L, McLaren-Douglas A, Blokker J, Jaksani S, Bartfeld S, Volckman R, van Sluis P, Li Vivian SW, Seepo S, Sekhar Pedamallu C, Cibulskis K, Carter Scott L, McKenna A, Lawrence Michael S, Lichtenstein L, Stewart C, Koster J, Versteeg R, van Oudenaarden A, Saez-Rodriguez J, Vries Robert GJ, Getz G, Wessels L, Stratton Michael R, McDermott U, Meyerson M, Garnett Mathew J, Clevers H (2015) Prospective Derivation of a Living Organoid Biobank of Colorectal Cancer Patients. Cell 161: 933–945 10.1016/j.cell.2015.03.053

32. Hicks WH, Bird CE, Gattie LC, Shami ME, Traylor JI, Shi DD, McBrayer SK, Abdullah KG (2022) Creation and Development of Patient-Derived Organoids for Therapeutic Screening in Solid Cancer. Current Stem Cell Reports 8: 107–117 doi:10.1007/s40778-022-00211-2

33. Chen CC, Li HW, Wang YL, Lee CC, Shen YC, Hsieh CY, Lin HL, Chen XX, Cho DY, Hsieh CL, Guo JH, Wei ST, Wang J, Wang SC (2022) Patient-derived tumor organoids as a platform of precision treatment for malignant brain tumors. Sci Rep 12: 16399 doi:10.1038/s41598-022-20487-y

34. Jacob F, Salinas RD, Zhang DY, Nguyen PTT, Schnoll JG, Wong SZH, Thokala R, Sheikh S, Saxena D, Prokop S, Liu DA, Qian X, Petrov D, Lucas T, Chen HI, Dorsey JF, Christian KM, Binder ZA, Nasrallah M, Brem S, O’Rourke DM, Ming GL, Song H (2020) A Patient-Derived Glioblastoma Organoid Model and Biobank Recapitulates Inter- and Intra-tumoral Heterogeneity. Cell 180: 188–204 e122 doi:10.1016/j.cell.2019.11.036

35. Ou L, Liu S, Wang H, Guo Y, Guan L, Shen L, Luo R, Elder DE, Huang AC, Karakousis G, Miura J, Mitchell T, Schuchter L, Amaravadi R, Flowers A, Mou H, Yi F, Guo W, Ko J, Chen Q, Tian B, Herlyn M, Xu X (2023) Patient-derived melanoma organoid models facilitate the assessment of immunotherapies. EBioMedicine 92: 104614 doi:10.1016/j.ebiom.2023.104614

36. El Shami M, Savani MR, Gattie LC, Smith B, Hicks WH, Rich JN, Richardson TE, McBrayer SK, Abdullah KG (2023) Human plasma-like medium facilitates metabolic tracing and enables upregulation of immune signaling pathways in glioblastoma explants. bioRxiv doi:10.1101/2023.05.29.542774

